# Immunogenicity and safety in pigs of PHH-1V, a SARS-CoV-2 RBD fusion heterodimer vaccine candidate

**DOI:** 10.1101/2023.01.19.524684

**Authors:** Alexandra Moros, Antoni Prenafeta, Antonio Barreiro, Eva Perozo, Alex Fernández, Manuel Cañete, Luis González, Carme Garriga, Edwards Pradenas, Silvia Marfil, Julià Blanco, Paula Cebollada Rica, Marta Sisteré-Oró, Andreas Meyerhans, Teresa Prat Cabañas, Ricard March, Laura Ferrer

## Abstract

The continuing high global incidence of COVID-19 and the undervaccinated status of billions of persons strongly motivate the development of a new generation of efficacious vaccines. We have developed an adjuvanted vaccine candidate, PHH-1V, based on a protein comprising the receptor binding domain (RBD) of the Beta variant of SARS-CoV-2 fused in tandem with the equivalent domain of the Alpha variant, with its immunogenicity, safety and efficacy previously demonstrated in mouse models. In the present study, we immunized pigs with different doses of PHH-1V in a prime-and-boost scheme showing PHH-1V to exhibit an excellent safety profile in pigs and to produce a solid RBD-specific humoral response with neutralising antibodies to 7 distinct SARS-CoV-2 variants of concern, with the induction of a significant IFNγ^+^ T-cell response. We conclude that PHH-1V is safe and elicits a robust immune response to SARS-CoV-2 in pigs, a large animal preclinical model.

## INTRODUCTION

Coronavirus disease-19 (COVID-19) was declared a pandemic by the World Health Organization (WHO) in March 2020 due to its widespread transmission, prompting a global effort to develop safe and effective vaccines against SARS-CoV-2, with more than 240 candidate vaccines currently under clinical development [1]. These include messenger RNA, protein-based and viral vector-based vaccines targeting the viral spike protein or its Receptor Binding Domain (RBD), and conventional inactivated vaccines targeting the entire virus [2]. Several vaccines have shown great efficacy in protecting from severe disease, and have been deployed and administered in an unprecedented global vaccination effort. However, the continued emergence of new viral variants with increasing immune escape and transmissibility characteristics compromises the intended control of the pandemic [3]. In addition, an unequal distribution of currently available vaccines is present, with over 80% of the population in low- and middle-income countries not having received any SARS-CoV-2 vaccine dose (as of July 2022) [1].

With these motivations, we have developed the adjuvanted PHH-1V vaccine candidate based on a fusion heterodimer protein containing, in tandem, the RBD of the B.1.351 (Beta) and B.1.1.7 (Alpha) SARS-CoV-2 variants at its N- and C-terminus, respectively. As a result, a key feature of PHH-1V distinguishing it from other adjuvanted protein-based subunit vaccines is the inclusion of sequences corresponding to two heterologous variants of concern (VOCs) as part of the same protein molecule. The safety, immunogenicity and efficacy of the PHH-1V vaccine candidate were previously tested in mouse models [4]. Recently, EMA’s human medicines committee ([CHMP](https://www.ema.europa.eu/en/glossary/chmp)) has recommended authorising the PHH-1V vaccine (Bimervax) as a booster in people aged 16 years and above who have been vaccinated with an mRNA COVID-19 vaccine.In the present study we have further investigated its safety and immunogenicity in pigs - a large animal model involving an immune system with physiologically relevant similarities to humans [5].

## MATERIALS AND METHODS

### PHH-1V vaccine

PHH-1V is a bivalent recombinant protein vaccine based on a heterodimer containing the receptor binding domain (RBD) of two SARS-CoV-2 variants, Beta and Alpha. The SARS-CoV-2 Wuhan-Hu-1 Spike RBD sequence corresponds to amino acids 319-541, according to the code P0DTC2 [6]. Based on this sequence, a dimer of RBDs was designed containing the mutations of the SARS-CoV-2 Beta variant (B.1.351) at its N-terminus and the mutation of the Alpha variant (B.1.1.7) at its C-terminus. The RBD heterodimer is expressed in CHO cells with stable expression (clone 57) and adjuvanted with SQBA, an oil-in-water emulsion based on squalene, in which the oil particles are dispersed to the aqueous phase. SQBA adjuvant contains per 0.5 mL dose: squalene (9.75 mg), polysorbate 80 (1.18 mg), sorbitan trioleate (1.18 mg), sodium citrate (0.66 mg), citric acid (0.04 mg), and water for injection.

### Animal study design

Large White x Landrace hybrid pigs were used as a model to assess immune responses to PHH-1V. In the first study (Study 1), three groups of six pigs (3 females and 3 males) were administered with three different doses of PHH-1V vaccine (Group A: 10 μg; Group B: 20 μg; Group C: 40 μg), and one group of four pigs (two females and two males) received PBS (Group D) **(Figure 1)**. In the second study (Study 2) for safety and cellular immune response analysis, ten male pigs were distributed into one group of six pigs administered 40 μg PHH-1V per dose (Group 1) and a second group of four pigs administered PBS (Group 2). In both studies, pigs received an intramuscular injection in a total volume of 0.5 mL of test product on days 0 (D0) and 21 (D21) **(Figure 1)**. Safety was assessed by monitoring general clinical signs (recorded daily) and rectal temperatures (recorded one day before each dosage, at vaccination, 4 and 6 hours after that, and daily for 5 days after the first injection and 2 days after the second injection) and local reactions (inflammation and nodules) at the injection site (observed daily) throughout the study. Blood samples for the different analyses were extracted from the external jugular vein of the animals.

**Figure 1.**
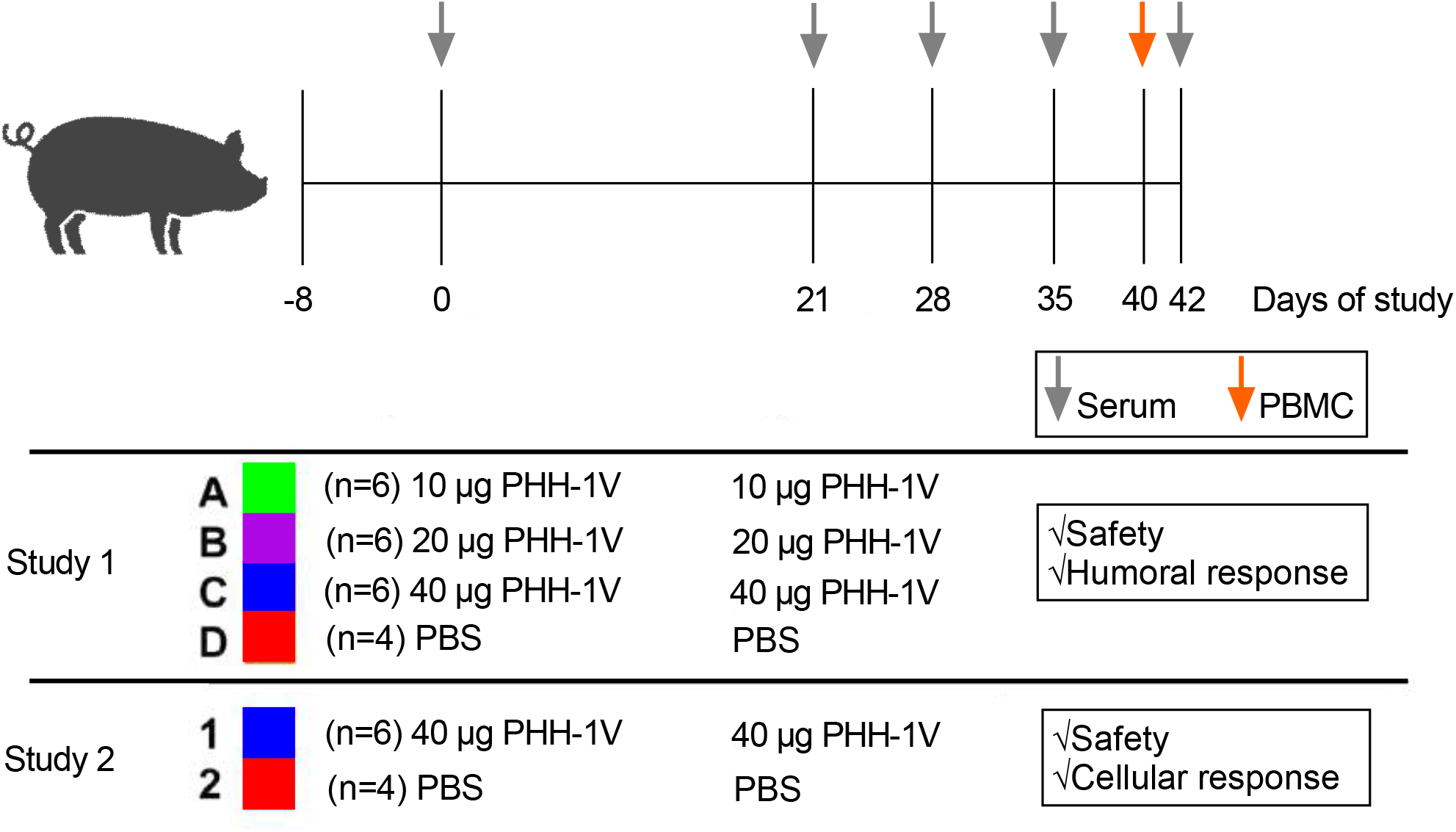
Experimental procedure schematics. Two-dose prime-and-boost intra-muscular immunization on days 0 and 21 (0.5 mL/dose), with a 42-day follow-up. Two independent studies were performed. Study 1: For safety and humoral response monitoring, three experimental groups (A-C) were inoculated (n = 6/group; 3 female and 3 male) with 10 μg, 20 μg or 40 μg of PHH-1V. A control group (D; n = 4; 2 female and 2 male) was inoculated with PBS. Study 2: For safety and cellular response monitoring, pigs of group 1 (n = 6; 6 male) were inoculated with 40 μg of PHH 1V. Control group 2 (n = 4; 4 male) was inoculated with PBS.

### *Ex vivo* assays

#### ELISA determinations for SARS-CoV-2 RBD-binding antibodies

MaxiSorp plates (Nunc) were coated with 100 μL/well of 1 μg/mL of SARS-CoV-2 RBD protein (Sino Biologicals) O/N at 4 ºC, then washed with 300 μL/well of CIVTEST solution (Laboratorios HIPRA) and blocked with 250 μL/well of StabilBlock Immunoassay Stabilizer (Surmodics) for 1 hour at room temperature. Wells were incubated with 100 μL of serial dilutions of the serum samples, then washed again, and after that, the bound total IgG specific antibodies were detected by 100 μL of peroxidase-conjugated AffiniPure Goat Anti-Swine IgG (H+L) (Jackson ImmunoResearch) incubated for 1 hour at 37 ºC. After washing, an incubation with 100 μL of TMB Substrate (Neogen) during 15 minutes at room temperature protected from light was done. Then 100 μL of TMB stop solution was added and measurement of absorbance at 450 nm in a Versamax instrument (Molecular Devices) was performed. The end-point titre of RBD-bound total IgG antibodies was established as the reciprocal of the last serum dilution that yielded three times the mean optical density of the negative control.

#### Pseudovirus-based neutralisation assays (PBNA)

Pseudoviruses were generated by cotransfection of Expi293F cells (Thermo Fisher Scientific) with the pNL4-3.Luc.R-.E-vector (NIH AIDS repository) and a pcDNA3.1(+) plasmid coding for SARS-CoV-2 S protein (GeneArt) with a deletion of the C-terminal 19 amino acids and human-codon optimised. The same procedure was followed to generate expression plasmids for the Alpha, Beta, Gamma, Delta, Delta plus and Omicron (BA.1) variants of the SARS-CoV-2 spike protein. Transfections were carried out using ExpiFectamine Reagent (Thermo Fisher Scientific) according to manufacturer instructions at an 8:1 ratio (HIV:Spike). After 48 hours of culture at 37 ºC and 8% CO_2_, supernatants were harvested, filtered through 0.45 μm filters, frozen, and subsequently titrated on HEK293T cells overexpressing human ACE-2 (Integral Molecular).

The neutralization assay has been previously validated in a large subset of samples with a replicative viral inhibition assay [7]. Neutralization assays were performed in duplicate in Nunc 96-well cell culture plates (Thermo Fisher Scientific), 200 TCID_50_ of pseudovirus were preincubated with three-fold serial dilutions (1/60–1/14,580) of heat-inactivated plasma samples for 1 hour at 37 ºC in a total volume of 150 μL. Then, 2×10^4^ HEK293T/hACE2 cells treated with DEAE-Dextran (Sigma-Aldrich) were added to each well. After 48 hour-incubation at 37 ºC and 5% CO_2_, luminiscence was determined using the EnSight Multimode Plate Reader and BriteLite Plus Luciferase reagent (PerkinElmer). Percentage neutralisation was calculated as (RLUmax–RLUexp)/(RLUmax–RLUmin)*100, with RLUexp being the experimental infectivity calculated from infected cells treated with each serum, RLUmax the maximal infectivity calculated from untreated infected cells, and RLUmin the minimal infectivity calculated from uninfected cells. IC_50_ values were calculated by fitting the neutralisation values and the log of plasma dilution to a 4-parameter equation using Prism 9.0.2 (GraphPad Software, USA).

#### RBD-induced IFNγ ELISpot assay

A total of 5×10^5^ peripheral blood mononuclear cells (PBMCs) per well in 100 μL/well were plated and *ex vivo* stimulated with other 100 μL/well of a 1:1 mixture of RBD-overlapping peptides from the SARS-CoV-2 B.1.1.7 and B.1.351 lineages (1 μg/mL per peptide, final concentration), with complete RPMI (negative control), or with 2.5 ng/mL PMA plus 250 ng/mL ionomycin (positive control; both from Sigma). After 18-20 hours incubation at 37 ºC, plates were developed using reagents and following procedures and incubation periods described in the pig IFNγ ELISpot Plus kit (3130-4HPW-10, Mabtech). Spots were counted under a dissection microscope (Leica GZ6). Data were expressed as the number of RBD-specific IFNγ-secreting cells per 1×10^6^ PBMCs.

### Statistical analysis

Statistical analyses and plots were generated using R (version 4.0.5) or GraphPad Prism (version 9). All plots depict individual data points for each animal, along with the sample mean and standard deviation. The exact number (n) used in each experiment is indicated in the caption below each figure. Data transformations were conducted to ensure a better distribution for appropriate model fitting: for the analysis of log-normal variables (IgG and neutralising antibodies) data was log_10_-transformed, while for the analysis of percentage variables (ELISpot and intracellular staining) data was arcsine-square root-transformed. Linear mixed effects models of repeated measures were fitted with the longitudinal data. In these models, the treatment group, the timepoint and the group-by-timepoint interaction were introduced as fixed effects and the animal ID as a random effect. Non-significant terms (at the 5% level) were sequentially removed from the models (backward elimination). Linear models were employed for the analysis of the temperature and humoral and cellular immunity data. For the temperature and IgG data, mixed effects models were used to account for the repeated measures design of the experiments, considering time and treatment group (and the two-way interaction, if supported by the model) as fixed factors and the animal identifier as random effect. For the analysis of the neutralising antibodies, a mixed effects model was used to account for the repeated measures design of the experiment, considering the treatment group and the SARS-CoV-2 variant as fixed factors and the animal identifier as a random effect. Also for neutralising antibodies, vaccinated groups were analysed with an ANOVA and the pooled data (groups A, B, C) was compared against the PBS group by means of a one-sample t-test (null hypothesis=1.31). On the other hand, generalised least squares models were fitted for the cellular immunity data to account for data heteroskedasticity. For all models, parameter estimation was performed using restricted maximum likelihood and assumptions were tested graphically.

Statistically significant differences between groups are indicated with a line on top of each group: ^**^ p<0.01; ^*^ p<0.05; ^+^ 0.05<p<0.1.

## RESULTS

### PHH-1V displays a good safety profile in pigs

Rectal temperatures showed no differences between groups (**Supplementary Figure 1**). Local reactions at the injection site or general clinical signs were not observed in any animal throughout the study. Furthermore, in the Study 1, the maximum individual increases after first and second administration were 0.59 ºC and 0.68 ºC, respectively, in both cases in group B. On the other hand, in the Study 2, the maximum individual temperature increases after first and second administration were 1.16 ºC and 0.43 ºC, respectively, in both cases in group 1.

### PHH-1V elicits a robust humoral response

After acclimation, pigs were immunized (day 0) and boosted (day 21) with three different doses of PHH-1V. Seven days after the second vaccination (day 28), the sera from all the animals were analysed by ELISA to determine the RBD-binding antibodies. Immunization induced significant levels of RBD-specific IgG antibodies in all vaccinated groups compared to the control group (**Figure 2A and Supplementary Table 1**). In all dosage groups, the antibody titres remained at stably elevated levels until the end of the study on day 42, with significant differences compared to control animals (**Figure 2A**).

**Figure 2.**
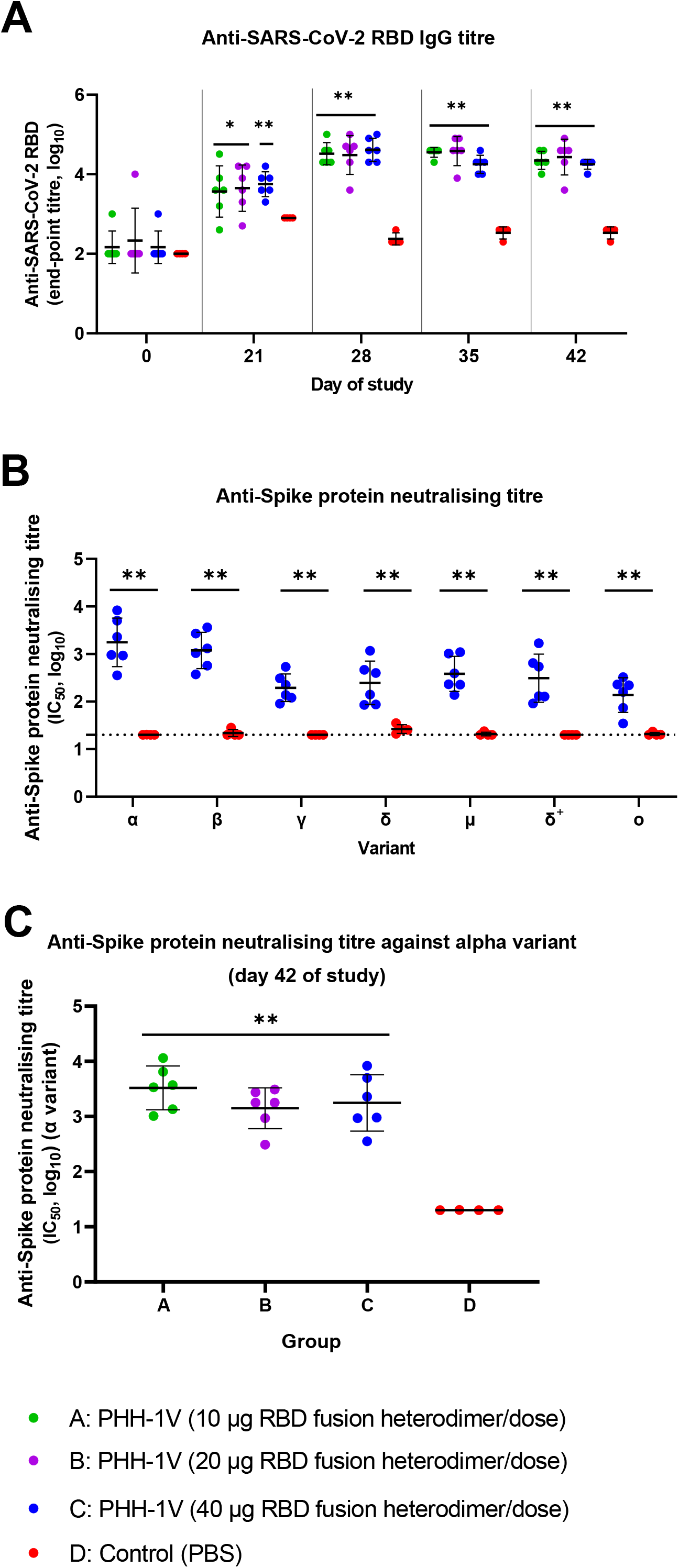
Vaccination of pigs with PHH-1V elicits SARS-CoV-2 RBD-specific humoral and neutralising antibody responses against SARS-CoV-2 variants of concern (VOCs). **A**. Longitudinal analysis of RBD-specific IgG determined in samples collected on days 0, 21, 28, 35 and 42. Group A: 10 μg PHH-1V RBD fusion heterodimer; Group B: 20 μg PHH-1V RBD fusion heterodimer; Group C: 40 μg PHH-1V RBD fusion heterodimer; D: Control PBS. Each data point represents an individual pig with bars denoting the mean ± standard deviation per group. IgG longitudinal data have been analysed by means of a linear mixed effects model of repeated measures with the log_10_-transformed titres. ^*^*p*<0.05; ^**^ <0.01, indicating statistically significance vs. Control group (D). **B**. Neutralisation activity by sera collected on day 42 after the first vaccine dose from Group C (40 μg PHH-1V) vs. Group D (PBS) against 7 different VOCs: α (B.1.1.7), β (B.1.351), γ (P.1), μ (B.1.621), δ (B.1.617.2), δ+ (B.1.617.2.1) and o (BA.1). Pseudovirus neutralisation assays were conducted, yielding titres expressed as log_10_ IC_50_. Each data point represents an individual pig with bars denoting the mean ± standard deviation per group. PBNA IC_50_ longitudinal data have been analysed by means of a linear mixed effects model of repeated measures with the log_10_-transformed titres. ^**^ *p*<0.01. Titre values are provided in **Table S2. C**. Neutralization activity by sera collected on day 42 after the first vaccine dose from Group A: 10 μg PHH-1V RBD fusion heterodimer; Group B: 20 μg PHH-1V RBD fusion heterodimer; Group C: 40 μg PHH-1V RBD fusion heterodimer; and D: Control PBS. Pseudovirus neutralisation assays were conducted, yielding titres expressed as log_10_ IC_50_. Each data point represents an individual pig with bars denoting the mean ± standard deviation per group. PBNA data in vaccinated groups were compared with an ANOVA and the pooled data (groups A, B, C) was compared against the PBS group by means of a one-sample t-test (null hypothesis =1.31).. Differences between vaccinated groups were not significant but the three groups were significantly different from the control group. ^**^ *p*<0.01 indicating statistically significance vs. Control group (D).

### PHH-1V elicits neutralising antibodies to all SARS-CoV-2 variants of concern

A prime-boost of PHH-1V (40 μg/dose) elicited significant titres of neutralising antibodies (nAbs) against all VOCs analysed by PBNA compared to non-vaccinated animals in sera collected on day 42 post-prime dose (**Figure 2B**). When compared to the Alpha and Beta variants, the neutralising titre of the sera was slightly reduced against Gamma, Mu, Delta and Omicron BA.1 variants (**Figure 2B and Supplementary Table 2**). No significant differences were observed between PHH-1V doses in the neutralising titres against Alpha variant on day 42 (**Figure 2C**).

### PHH-1V elicits a robust cellular immune response

To determine the cellular immune response (Study 2), peripheral blood mononuclear cells (PBMCs) were isolated on day 40 and tested by IFNγ ELISpot *ex vivo* for their reactivity towards a pool of RBD peptides representing Alpha and Beta SARS-CoV-2 variant epitopes. The PBMCs from vaccinated pigs (40 μg RBD fusion heterodimer/dose) showed a significant higher percentage of IFNγ^+^ T-cells than in control animals after *in vitro* re-stimulation (**Figure 3A**).

**Figure 3.**
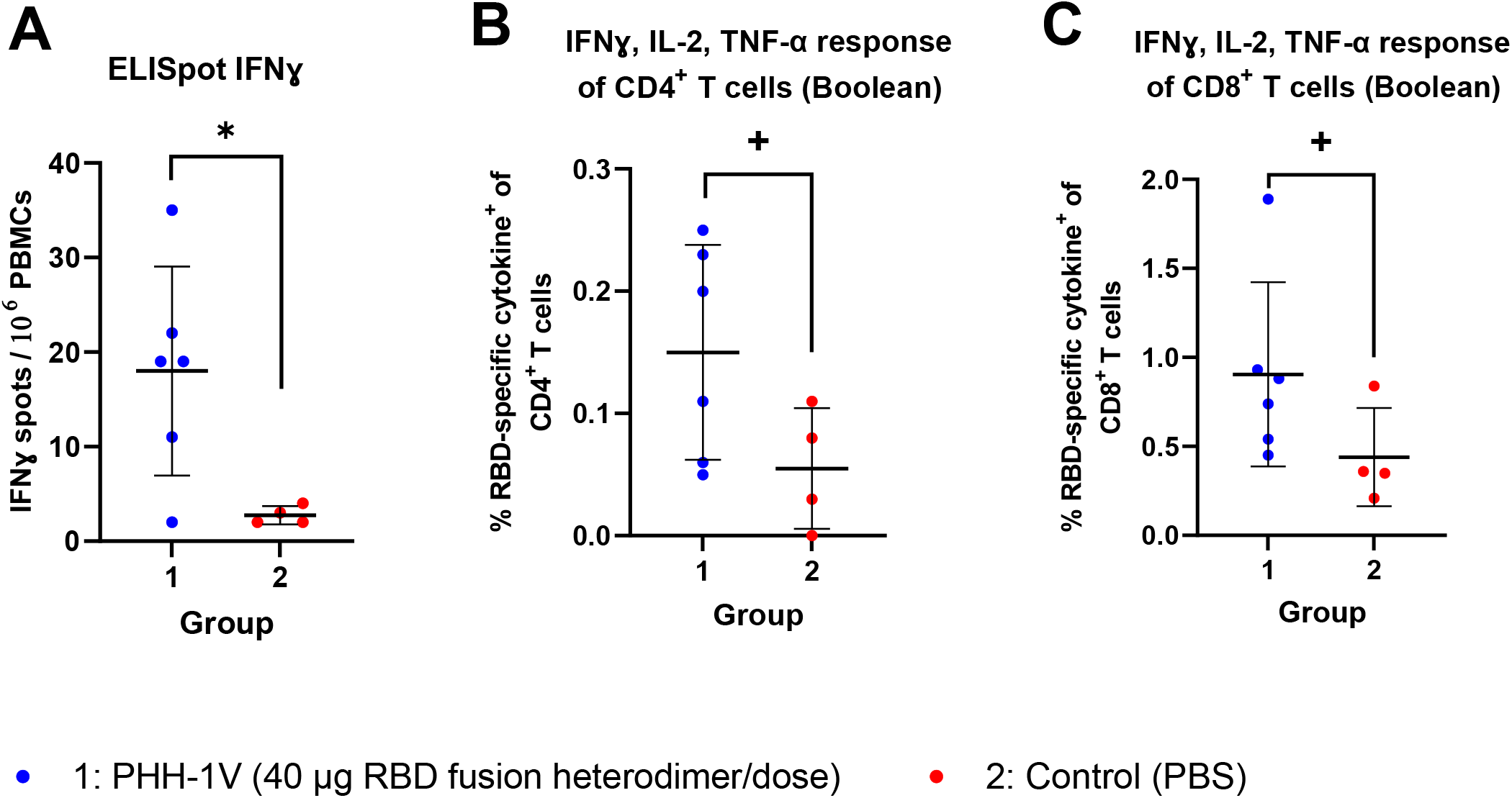
Vaccination of pigs with PHH-1V elicits cellular response. **A**. Induction of IFNγ^+^ T-cells determined by ELISpot assays in PBMCs from vaccinated pigs, sampled on day 40 and stimulated *ex vivo* with an RBD peptide pool representing Alpha and Beta SARS-CoV-2 variant epitopes. Group 1: 40 μg PHH-1V RBD fusion heterodimer; Group 2: Control PBS. Each data point represents an individual pig with bars denoting the mean ± standard deviation per group. ELISpot data have been analysed by means of a generalised least squares model with the arcsine-square root-transformed proportion of IFNγ -positive cells ^*^ p<0.05. **B-C**. CD4^+^ (**B)** and CD8^+^ **(C)** T cells expressing IFNγ and/or IL-2 and/or TNF-α was measured by intracellular cytokine staining (ICS) after the stimulation of PMBCs with a pool of peptides from SARS-CoV-2 RBD of Alpha and Beta variants. Cytokine expression of PBMCs stimulated with medium was considered as background and subtracted from the responses measured from the RBD peptide pool for each individual pig. Group 1: PHH-1V vaccine 40 μg RBD fusion heterodimer/dose; Group 2: control group (PBS). Each data point represents an individual pig with bars denoting the mean ± standard deviation per group. ICS data have been analysed by means of a generalised least squares model with the arcsine-square root-transformed proportion of positive cells ^+^ 0.05<*p*<0.1.

In addition, in PHH-1V immunised pigs a tendency was observed in the activation antigen-specific Th1 response of CD4^+^ and CD8^+^ T cells expressing IFNγ and/or IL-2 and/or TNF-α after *in vitro* re-stimulation **(Figures 3B and 3C)**. Supplementary Figure 3 shows individual frequencies of cytokine expressing T cells.

## DISCUSSION

An unprecedented effort has enabled the development of a substantial number of vaccines and several therapies against SARS-CoV-2, a novel virus that has caused an enormous impact on lives [1]. Many vaccine candidates have been designed, using broadly diverse platforms and technologies, which has afforded the production of the large numbers of doses required for mass vaccination programs. Despite the phenomenal success and effectiveness of several of these vaccines, the continuing pandemic, the insufficient distribution of vaccines to many developing countries, and the emergence of new immune escape variants endowed with increasing transmissibility [8], must be urgently addressed with an end to the pandemic as a goal - including the development of a new generation of affordable vaccines featuring higher efficacy against emerging variants and improved safety [3].

In the present study we have examined the safety and immunogenic potential of the RBD subunit-based vaccine candidate, PHH-1V, in pigs which is a large animal presenting an immune system with significant similarities to humans [5], and a useful pre-clinical model to for assessing the immunogenicity of upcoming SARS-CoV-2 vaccine candidates [9]. We have found that two vaccination doses are safe and induce high and specific humoral and cellular immune responses. As expected, PHH-1V elicited strong RBD-binding IgG antibodies after two doses. Notably, it also generated significant neutralisation activity, determined by pseudovirus-based assays, against all tested VOCs (Alpha, Beta, Gamma, Delta and Omicron BA.1). The results of ELISpot assaying showed that 40 μg of PHH-1V induces significantly higher percentage of T-cells expressing IFNγ than mock-vaccinated pigs, being this dose a candidate to induce both humoral and cellular responses.

Neutralising antibody levels correlate well with vaccine protective efficacy [10]. As the RBD bears immunodominant neutralisation epitopes [11], a vaccine consisting of RBD, rather than a full-length S protein, is expected to induce more effectively RBD-focused antibodies [12, 13]. A second crucial feature of PHH-1V is the simultaneous presence in the same protein of RBDs for two heterologous VOCs, Beta and Alpha, in a two-domain tandem fusion configuration [14]. We have found that PHH-1V induces neutralising antibodies that are cross-reactive against the majority VOCs with the strongest presence to date, including the highly immunoevasive Beta, Gamma and Omicron BA.1 variants, after two doses. The presence of known immune escape mutations on PHH-1V, such as K417N, E484K or N501Y [15, 16], may be partially responsible for the neutralisation cross-reactivity observed against variants sharing these mutations, such as Alpha (N501Y), Beta (K417N, E484K, N501Y), Gamma (E484K, N501Y), Mu (E484K) or Omicron (K417N, N501Y), although epitopes not containing such immune escape mutations must play a role in eliciting neutralising antibodies against the Delta SARS-CoV-2 variant [17-19].

A limitation of the present study is that a challenging analysis after the PHH-1V vaccine is not feasible, as the pig model is not susceptible to SARS-CoV-2 [20]. Nevertheless, our previous challenge studies in mice models and non-human primates have demonstrated efficacy, safety and protection from the viral challenge of PHH-1V [4]. Future studies should address additional relevant aspects, such as the immune response to booster doses or the induction of B and T cell memory.

## CONCLUSIONS

In conclusion, PHH-1V exhibited an excellent safety profile and a strong and specific immune response in pigs, showing a broadly cross-reactive neutralisation activity against heterologous VOCs, and IFNγ^+^ T-cells activation. These results and those observed in other pre-clinical models, support the implementation of the PHH-1V vaccine as a primary or booster vaccination schedule in humans to prevent the consequences of SARS-CoV-2 infection. Further Phase IIb and III clinical trials are ongoing to support the potential of this candidate vaccine as a heterologous booster (NCT05142553, NCT05246137) [21, 22].

## ACKNOWLEDGEMENTS

Anna Moya and Mireia Muntada for the ELISA analysis; Clara Panosa and Ester Puigvert for her assistance in the production of the vaccine antigen; Glòria Pujol and Eduard Fossas for their assistance in review of the manuscript; and Alicia Subtil-Rodríguez from Dynamic S.L.U. (Evidenze Clinical Research, Madrid, Spain) for providing medical writing support during the preparation of this paper funded by Hipra Scientific, S.L.U.

