## Supplementary Material for "Immunogenicity and safety in pigs of PHH-1V, a SARS-CoV-2 RBD fusion heterodimer vaccine candidate"

### SUPPLEMENTARY DATA

#### SUPPLEMENTARY MATERIALS AND METHODS

##### Flow cytometry

Isolated pig PBMCs were seeded at $2x10^7$ cells/mL and stimulated for 5 hours with a 1:1 admixture of peptide libraries (PepMix™) from SARS-CoV-2 lineages B.1.1.7 and B.1.351 covering the respective RBDs. For positive and negative controls, PMA (Sigma) + ionomycin (Sigma) and cRPMI were used, respectively. Brefeldin A (10 µg/mL; BFA, Sigma) was added to block cytokine secretion for the last three hours. For T cell surface markers and cytokine characterisation, cells were labelled with Zombie NIR viability stain (BioLegend) and CD4-PerCP-Cy5.5 (74-12-4, BD Bioscience) or CD8-β-FITC (PPT23, Bio-Rad) fluorescence-labelled monoclonal antibodies. Cells were later fixed with 2% formaldehyde and stained with IFNγ-AF647 (CC302, Bio-Rad), TNF-α-BV421 (Mab11, BioLegend), IL-2 biotin (ASC0829 and clone A150D8H10, Thermofisher) or IL-4 BV605 (MP4-25D2, BioLegend) and streptavidin PE-Cy7 conjugate (eBioscience, ThermoFisher Scientific). Detection and quantitation were performed on an Aurora (Cytek) flow cytometer. To quantify the specific Th1 response from $CD4^+$ and $CD8^+$ T cells, gates for TNF-α and/or IL-2 and/or $IFNγ^+$ $CD4^+$ or $CD8^+$ T cells were combined and analysed by the Boolean tool of the FlowJo software.

#### SUPPLEMENTARY FIGURES LEGENDS

**Supplementary Figure 1: Average temperature increase (ºC) after vaccine administration of study 1 (A) and study 2 (B).** Data is presented as a box plot: the median marks the mid-point on the data and is shown by the line that divides the box into two parts; inferior and superior limit of the box represent the first and third quartile, respectively; the upper and lower whiskers represent the minimum and maximum values of the distribution. Linear mixed effects models of repeated measures were fitted with the temperature longitudinal data. In these models, the treatment group, the timepoint and the group-by-timepoint interaction were introduced as fixed effects and the animal ID as a random effect. Non-significant terms (at the 5% level) were sequentially removed from the models (backward elimination). Eventually, only the timepoint factor was significant, hence suggesting that there is not enough evidence to support differences between groups in the temperature at any timepoint.

**Supplementary Figure 2: Identification of IFN-γ, IL-2, TNF-α and IL-4 secreting $CD4^+$ T cells and IFN-γ, IL-2, TNF-α and IL-4 secreting $CD8^+$ T cells from pig PBMCs using flow cytometry.**

Representative gating strategy for PBMC from non-immunized pigs: CD4+ and CD8+ T cells were selected as alive and lineage negative (γδ-T).

**Supplementary Figure 3: Cellular response after the stimulation with a pool of peptides from SARS-CoV-2 RBD of Alpha and Beta variants.** $CD4^+$ (**B**) and $CD8^+$ **(C)** T-cell response was measured by intracellular cytokine staining (ICS) after the stimulation of PMBCs with a pool of peptides from SARS-CoV-2 RBD of Alpha and Beta variants. Cytokine expression (IFN-γ and/or TNF-α and/or IL-2 and/or IL-4) of PBMCs stimulated with medium was considered as background and subtracted from the responses measured from the RBD peptide pool for each individual pig. Group 1 (blue): PHH-1V vaccine 40 µg RBD fusion heterodimer/dose; Group 2 (red): control group (PBS). Each data point represents an individual pig with bars denoting the mean ± standard deviation per group. ICS data have been analysed by means of a generalised least squares model with the arcsine-square root-transformed proportion of positive cells $^+$ $0.05<p<0.1$.

#### SUPPLEMENTARY TABLES

**SUPPLEMENTARY TABLE 1: Levels of RBD-specific antibodies by animal group and day of sample collection.**

| | | IgG titre ($log_{10}$ end-point) | | | | |
|---|---|---|---|---|---|---|
| **Group** | **Sex** | **Day 0** | **Day 21** | **Day 28** | **Day 35** | **Day 42** |
| A | F | 2.00 | 2.60 | 4.30 | 4.30 | 4.00 |
| A | F | 2.00 | 3.60 | 4.60 | 4.60 | 4.30 |
| A | F | 3.00 | 4.51 | 5.00 | 4.60 | 4.60 |
| A | M | 2.00 | 3.60 | 4.30 | 4.60 | 4.30 |
| A | M | 2.00 | 3.90 | 4.30 | 4.60 | 4.30 |
| A | M | 2.00 | 3.20 | 4.60 | 4.60 | 4.60 |
| B | F | 2.00 | 3.90 | 5.00 | 4.90 | 4.90 |
| B | F | 2.00 | 3.30 | 3.60 | 3.90 | 3.60 |
| B | F | 2.00 | 4.20 | 4.60 | 4.60 | 4.60 |
| B | M | 2.00 | 3.60 | 4.70 | 4.90 | 4.60 |
| B | M | 4.00 | 4.20 | 4.70 | 4.60 | 4.60 |
| B | M | 2.00 | 2.70 | 4.30 | 4.60 | 4.30 |
| C | F | 3.00 | 3.90 | 5.00 | 4.00 | 4.30 |
| C | F | 2.00 | 3.90 | 4.90 | 4.00 | 4.30 |
| C | F | 2.00 | 4.20 | 4.60 | 4.60 | 4.30 |
| C | M | 2.00 | 3.60 | 4.30 | 4.30 | 4.00 |
| C | M | 2.00 | 3.60 | 4.60 | 4.30 | 4.30 |
| C | M | 2.00 | 3.30 | 4.30 | 4.30 | 4.30 |
| D | F | 2.00 | 3.51 | 2.30 | 2.30 | 2.30 |
| D | F | 2.00 | 3.51 | 2.60 | 2.60 | 2.60 |
| D | M | 2.00 | 3.20 | 2.30 | 2.60 | 2.60 |
| D | M | 2.00 | 2.90 | 2.30 | 2.60 | 2.60 |

**Abbreviations:** F = female; M = male.

**SUPPLEMENTARY TABLE 2: Titres of neutralising antibodies against all variants of concern.**

| | | | $\log_{10} IC_{50}$ | | | | | | |
|---|---|---|---|---|---|---|---|---|---|
| **Group** | **Sex** | **Day** | **α** | **β** | **γ** | **δ** | **μ** | **δ+** | **o** |
| C | F | 42 | 3.92 | 3.56 | 2.73 | 3.07 | 3.01 | 3.23 | 2.34 |
| C | F | 42 | 3.36 | 3.43 | 2.49 | 2.67 | 3.04 | 2.81 | 2.51 |
| C | F | 42 | 3.70 | 3.12 | 2.14 | 2.60 | 2.59 | 2.73 | 2.36 |
| C | M | 42 | 2.55 | 2.57 | 1.78 | 1.93 | 2.14 | 2.11 | 1.78 |
| C | M | 42 | 2.98 | 2.75 | 1.96 | 1.94 | 2.36 | 1.96 | 1.78 |
| C | M | 42 | 2.97 | 3.02 | 2.08 | 2.15 | 2.36 | 2.11 | 1.78 |
| D | F | 42 | 1.30 | 1.30 | 1.30 | 1.32 | 1.30 | 1.30 | 1.37 |
| D | F | 42 | 1.30 | 1.30 | 1.30 | 1.41 | 1.38 | 1.30 | 1.31 |
| D | M | 42 | 1.31 | 1.45 | 1.30 | 1.55 | 1.30 | 1.30 | 1.30 |
| D | M | 42 | 1.30 | 1.30 | 1.30 | 1.41 | 1.30 | 1.30 | 1.30 |

**Abbreviations:** F = female; M = male.
