## Supplementary Figures for "Immunogenicity and safety in pigs of PHH-1V, a SARS-CoV-2 RBD fusion heterodimer vaccine candidate"

A

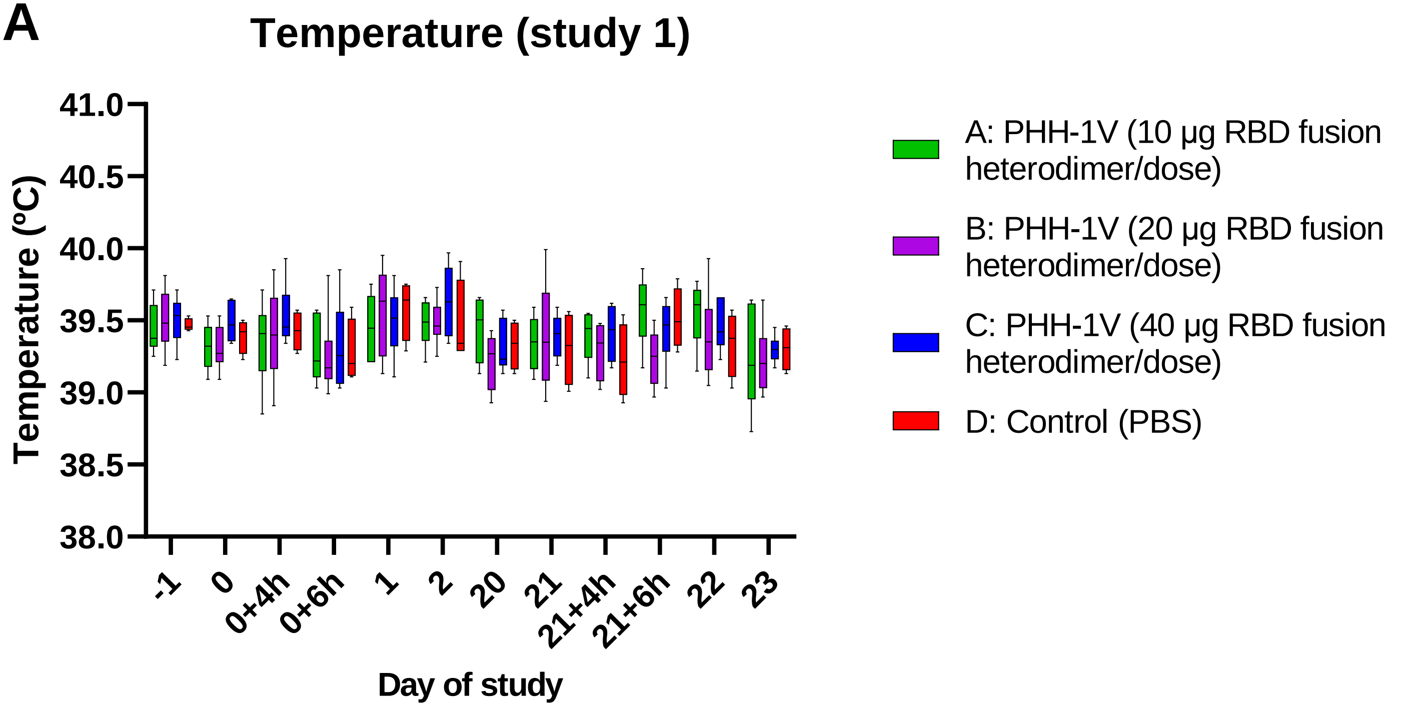


B

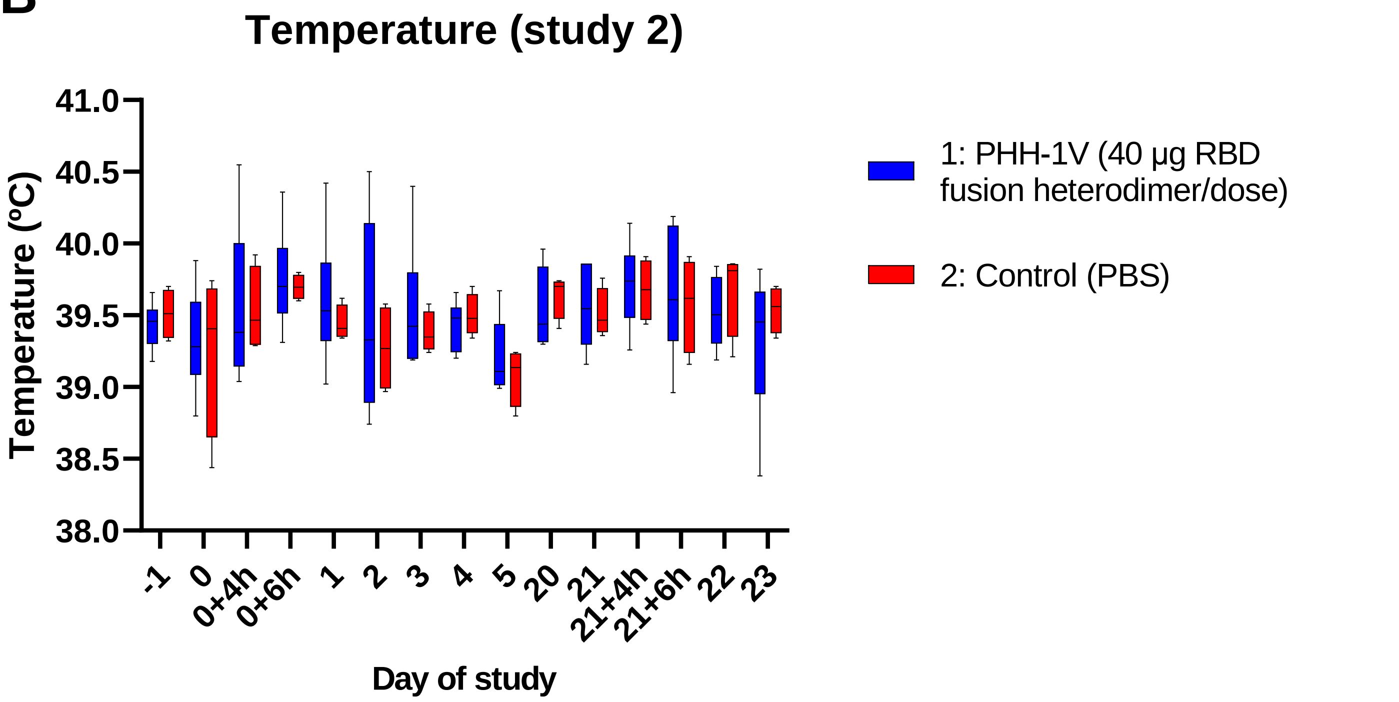


Supplementary 1

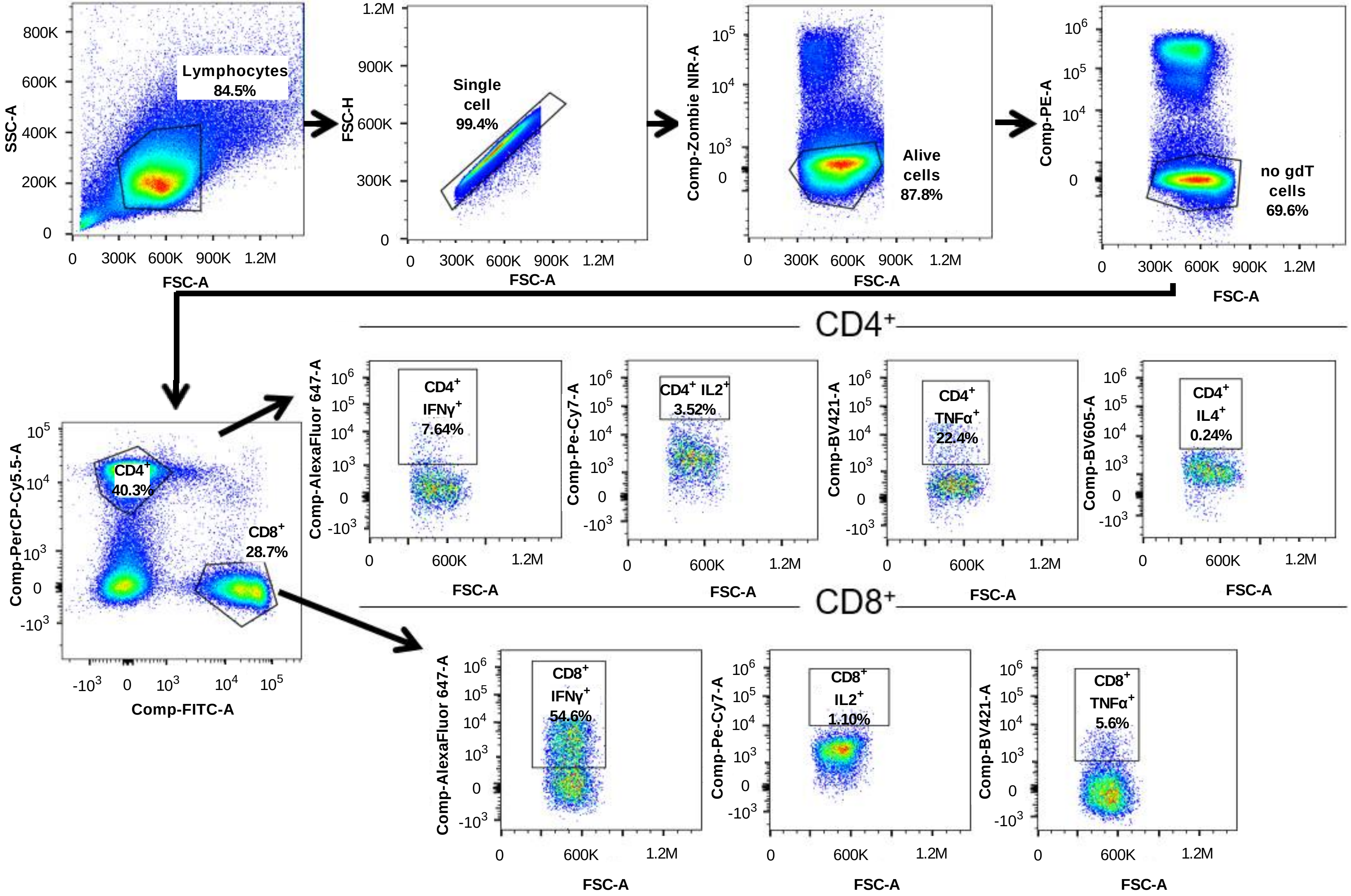


**Supplementary 2**

### IFNɣ, IL-2, TNF-α response of CD4+ T cells

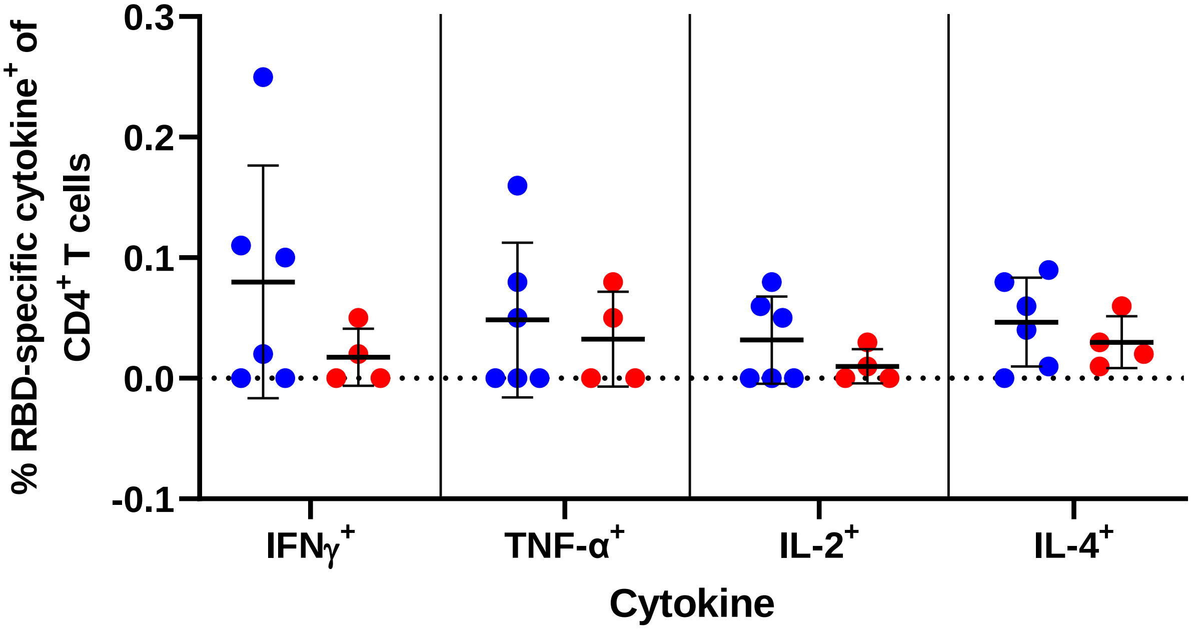


### IFNɣ, IL-2, TNF-α response of CD8+ T cells

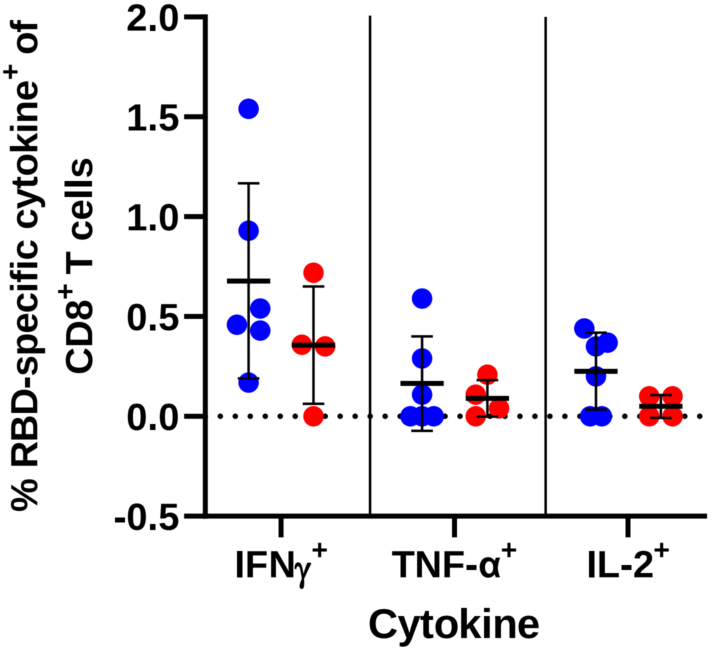


• 1: PHH-1V (40 µg RBD fusion heterodimer/dose) • 2: Control (PBS)

**Supplementary 3**
